# Setdb1 and Atf7IP form a hetero-trimeric complex that blocks Setdb1 nuclear export

**DOI:** 10.1101/2024.12.23.630145

**Authors:** Leena Kariapper, Ila A. Marathe, Ashley B. Niesman, Kelly Suino-Powell, Yuh Min Chook, Vicki H. Wysocki, Evan J. Worden

**Author notes:** Correspondence (EJW).

## Abstract

Histone H3K9 methylation (H3K9me) by Setdb1 silences retrotransposons (rTE) by sequestering them in constitutive heterochromatin. Atf7IP is a constitutive binding partner of Setdb1 and is responsible for Setdb1 nuclear localization, activation and chromatin recruitment. However, structural details of the Setdb1/Atf7IP interaction have not been evaluated. We used Alphafold2 predictions and biochemical reconstitutions to show that one copy of Setdb1 and two copies of Atf7IP form a hetero-trimeric complex *in vitro* and *in cells*. We also find that Atf7IP self-associates, forming multimeric complexes that are resolved upon Setdb1 binding. Setdb1 binds to Atf7IP through coiled coil interactions that include both Setdb1 nuclear export signals (NES). Atf7IP directly competes with CRM1 to bind the Setdb1 NES motifs, explaining how Atf7IP prevents CRM1-mediated nuclear export of Setdb1. Setdb1 also forms hetero-trimeric complexes with the Atf7IP paralog Atf7IP2 and we show that Setdb1 can form mixed heterotrimers comprising one copy of each Setdb1, Atf7IP and Atf7IP2. Atf7IP and Atf7IP2 are co-expressed in many tissues suggesting that heterotrimers with different compositions of Atf7IP and Atf7IP2 may differentially regulate H3K9me by fine-tuning Setdb1 localization and activity.

## Introduction

Retrotransposons (rTE) make up almost 45% of the human genomic DNA and have expanded to this extent due to the “copy-paste” mechanism of retro-transposition that allows rTEs to “copy” themselves through an RNA intermediate before reinserting into a different part of the genome^1, 2^. These ancient genetic elements have played a major role in shaping our genome and contribute to our genetic diversity, but also continue to pose serious risks to genome integrity and human health. While the huge majority of rTEs in humans cannot reinsert due to accumulated mutations^3^, expression of rTE RNA itself (without reinsertion) can trigger various negative effects ranging from inflammation to cell death^4–7^. Therefore, the cell tightly regulates rTE transcription by marking rTE chromatin with histone H3K9 di-, and tri-methylation (H3K9me2-3)^5, 8^ and DNA CpG methylation (mCpG)^9–11^. These epigenetic modifications silence expression of rTEs by sequestering them into constitutive heterochromatin. The H3K9 methyltransferase Setdb1 is responsible for most H3K9 methylation at retrotransposon sequences and is a critical component of the rTE silencing machinery^12^. Setdb1 loss causes global rTE activation in ES cells and cell type specific activation of certain rTEs in somatic tissues^13^. Dysregulated Setdb1 activity can also play a role in several human diseases. Setdb1 deficiency causes constitutive expression of rTEs in inflammatory bowel disease^14^ and overexpression of Setdb1 in cancers improves immune evasion by restricting expression of rTE-encoded proteins^15^. Therefore, Setdb1 is critical for regulating rTE expression during development and disease.

Because of its critical role in rTE silencing, the activity of Setdb1 is tightly regulated through its catalytic activation and recruitment to rTEs. The activity of Setdb1 is repressed in isolation, but Atf7IP binding directly stimulates its catalytic activity and prevents its nuclear export or degradation by the proteasome^16^. Atf7IP-dependent Setdb1 nuclear localization also stimulates mono-ubiquitination of the Setdb1 catalytic domain by the nuclear E2 enzyme Ube2e1^17, 18^. Mono-ubiquitination of Setdb1 further stimulates its catalytic activity, so the fully active catalytic module of Setdb1 is thought to be the mono-ubiquitinated Setdb1/Atf7IP complex^18^. However, because Atf7IP binding and Setdb1 mono-ubiquitination are tightly linked in the cell through nuclear localization of Setdb1, the relative contribution of each process to Setdb1 catalytic activation is not well understood. The Setdb1/Atf7IP catalytic module is then recruited to rTEs by various silencing complexes including KRAB zinc finger proteins^19^, the HUSH complex^20^, the Piwi silencing complex^21^, KDM5B^22^ and the m6A reader protein METTL3^23^. Importantly, the ubiquitinated Setdb1/Atf7IP module is primarily recruited to rTE chromatin for silencing through interactions with Atf7IP^18, 24^. Therefore, the interaction between Atf7IP and Setdb1 is critical for all aspects of Setdb1 dependent silencing, explaining why Atf7IP deletions phenocopy deletions of Setdb1^20^.

Although the interaction between Setdb1 and Atf7IP is required for rTE silencing, very little is known about how Setdb1 binds to Atf7IP. Previous work has shown that the N-terminal region of Setdb1 binds to Atf7IP and that a central fragment of Atf7IP interacts with Setdb1^18, 25^(**Fig 1a**), but it is not known if these two regions interact directly with each other. In addition, the N-terminal region of Setdb1 contains nuclear export signals (NES) that drive export of free Setdb1 out of the nucleus^25, 26^. While Atf7IP binding is required to prevent export of Setdb1 it is not clear *how* Atf7IP binding prevents export of Setdb1. Similarly, Atf7IP2, a meiosis-specific paralog of Atf7IP also regulates Setdb1’s nuclear localization, prevents its nuclear export and recruits it to specific chromatin loci in germ cells^27^. However, it is not clear if Atf7IP2 and Atf7IP bind to Setdb1 in a similar manner.

**Figure 1:**
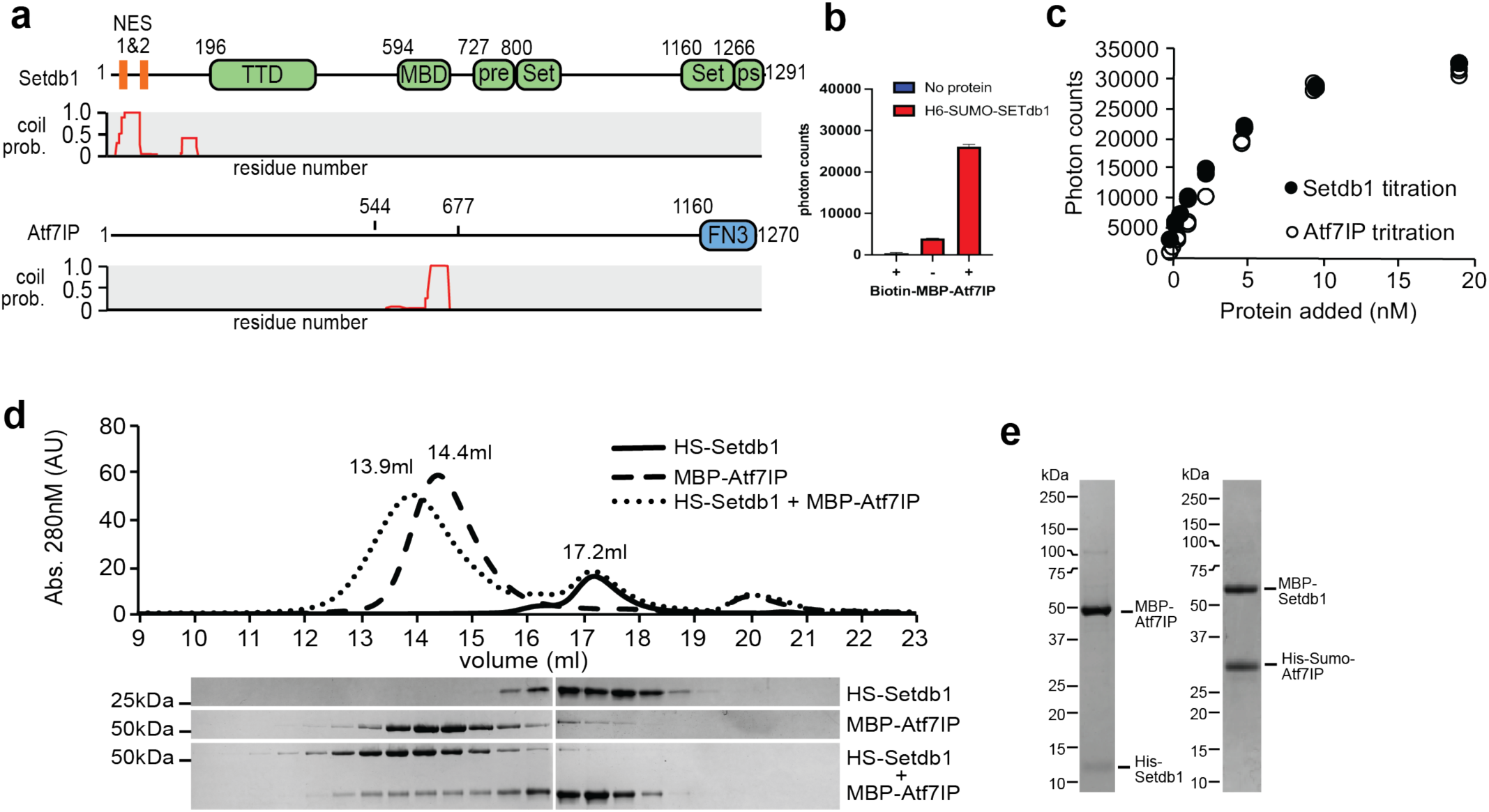
Setdb1 and Atf7IP interact through their coiled coil domains. a) Domain architecture of Setdb1 and Atf7IP (top) with the corresponding pcoils prediction for each protein (bottom). TTD = Triple Tudor Domain, MBD = methyl binding domain, pre = pre-set domain, Set = set domain, ps =post set domain, FN3 = fibronectin type 3 domain. b) Alphascreen binding assay using Ni^2+^ and streptavidin beads showing interaction his-sumo-Setdb1(2-115) and biotin-MBP-Atf7IP(574-666). c) Alphascreen assay as shown in b, but with increasing amounts of his-sumo-Setdb1 (closed circles) or increasing amounts of biotin-MBP-Atf7IP(574-666) (open circles). d) Superose 6 size exclusion chromatogram (top) showing elution of his-sumo (HS)-Setdb1(2-115), MBP- Atf7IP(574-666) and the HS-Setdb1(2-115)/Atf7IP(574-666) complex. SDS-PAGE gels (bottom) of the elution fractions from the size exclusion run. e) SDS-page gel bands of purified complexes 6Xhis-Setdb1(11-104)/MBP-Atf7IP(585-666) and MBP-Setdb1(2-115)/6Xhis-SUMO-Atf7IP(574-666).

Here we show that the N-terminal region of Setdb1 interacts with the central fragment of Atf7IP to form a minimal Setdb1/Atf7IP complex. Alphafold2 models of this complex reveal an extended, highly helical interaction with coiled coil elements. Surprisingly, we show that the Setdb1/Atf7IP complex is heterotrimeric, containing two copies of Atf7IP with one copy of Setdb1 and that Atf7IP forms homo-oligomers in the absence of Setdb1. NES motifs in the N-terminal region of Setdb1 are blocked within the Setdb1/Atf7IP heterotrimer and Atf7IP directly competes with the Chromosome Region Maintenance 1 (CRM1) nuclear export protein for Setdb1 binding, explaining its role in nuclear retention of Setdb1. Finally, we find that Atf7IP2 binds to Setdb1 with a similar heterotrimeric stoichiometry and that Atf7IP2 readily forms mixed trimers with Atf7IP and Setdb1 *in vitro*, suggesting that Setdb1 recruitment may be regulated by tuning the composition of Setdb1/Atf7IP/Atf7IP2 heterotrimers.

## Results

### Setdb1 and Atf7IP interact using coiled coil domains

Prior studies have shown that Atf7IP interacts with the first 109 amino acids of Setdb1(1-109) and that Setdb1 interacts the central region of Atf7IP spanning residues 562-817^16^ (**Fig. 1a**). We analyzed the sequences of Setdb1 and Atf7IP using the pcoils server^28^ and observed that these regions have a high probability to form coiled coils (**Fig. 1a**), suggesting that these coiled coil regions may directly interact with each other. Therefore, to determine if Setdb1 and Atf7IP form a complex mediated by these regions with high coiled coil propensity, we individually purified His-sumo-Setdb1(2-115) and biotin-labeled MBP-Atf7IP(574-666) and measured binding between both proteins using Alphascreen^29^, a bead-based assay of molecular interactions. We observed high Alphascreen levels when both His-sumo-Setdb1(2-115) and biotin-MBP-Atf7IP(574-666) were included in the binding reaction but not in reactions including only the individual proteins (**Fig 1b**). In addition, titration of His-sumo-Setdb1(2-115) into biotin-MBP-Atf7IP(574-666) produced a concentration-dependent increase of Alphascreen signal that was recapitulated when the titration was reversed (**Fig. 1c**). To confirm our Alphascreen results, we measured the interaction between His-sumo-Setdb1(2-115) and MBP-Atf7IP(574-666) by size exclusion chromatography (**Fig. 1d**). In isolation, Setdb1 and Atf7IP elute as single peaks at 17.2ml and 14.4ml, respectively. However, when a 2-fold excess Setdb1 was mixed with Atf7IP prior to injection, a fraction of Setdb1 coeluted with Atf7IP within a new peak centered at 13.9ml (**Fig. 1d**) suggesting that Setdb1 and Atf7IP directly interact. To confirm that the tags on Setdb1 and Atf7IP do not affect binding, we co-expressed the Setdb1 and Atf7IP truncations in E. coli with different tags and purified the resulting complexes (**Fig. 1e**). In all cases, both proteins stably co-purified, indicating that the identity of the fusion tag does not affect binding between these proteins. Together, these data indicate that the N-terminal region of Setdb1 stably binds to the central region of Atf7IP, potentially through coiled coil interactions.

### The Setdb1/Atf7IP complex is a heterotrimer

To understand how Setdb1 and Atf7IP interact, we sought to determine a crystal structure of the Setdb1/Atf7IP complex, however we were not able to grow crystals. Therefore, we modeled the Setdb1(2-115)/Atf7IP(574-666) complex using Alphafold2 multimer (**Fig. 2a,b**). Recent studies using Alphafold2 have shown that many proteins form oligomeric structures with various stoichiometries^2, 30^. Therefore, we modeled the Setdb1/Atf7IP complex by predicting structures using various stoichiometries of Setdb1 and Atf7IP (**Fig. S1**). Surprisingly, Alphafold2 generated two high-confidence models for the complex between Setdb1 and Atf7IP that differed in the number of Atf7IP molecules (**Fig 2a,b**). Alphafold2 predicted a 1:1 Setdb1:Atf7IP complex with very high confidence, as evident from the high pLDDT and PAE scores for this model (**Figs. S2a-b**). Alphafold2 also predicted a 1:2 Setdb1:Atf7IP complex, although this model was predicted with less confidence based on its lower pLDDT and PAE scores (**Figs. S2c-d**). In the 1:1 model, the three N-terminal helices of Setdb1 wrap around the two helices of Atf7IP spanning resides 574-666 (Fig. 2a). The second copy of Atf7IP in the 1:2 complex is split into three helices that bend around the 1:1 complex to contact the first copy of Atf7IP and Setdb1 (**Fig. 2b**). The central and C-terminal regions of the 1:2 complex have clear coiled coil character, agreeing with the prediction by the pcoils server, and the central region is almost perfectly superimposable with an idealized three-helix coiled coil^31^ (PDB:4dzl) (**Fig. S2f**). The 1:1 complex, while still resembling a coiled coil, does not superimpose as well to an idealized two helix coil (PDB: 4dzm)^31^ (**Fig. S2e**). The overall conformation of the 1:1 Setdb1:Atf7IP complex is preserved in the 1:2 complex and the two models are superimposable (RMSD 0.86Å) with the primary difference being the presence of a second Atf7IP molecule (**Fig. S2g**). Together these Alphafold2 models suggest that Setdb1/Atf7IP binding occurs through an alpha helical interface that is partially composed of coiled coil interactions. However, the precise stoichiometry of the complex cannot be conclusively determined using only Alphafold2 modeling.

**Figure 2:**
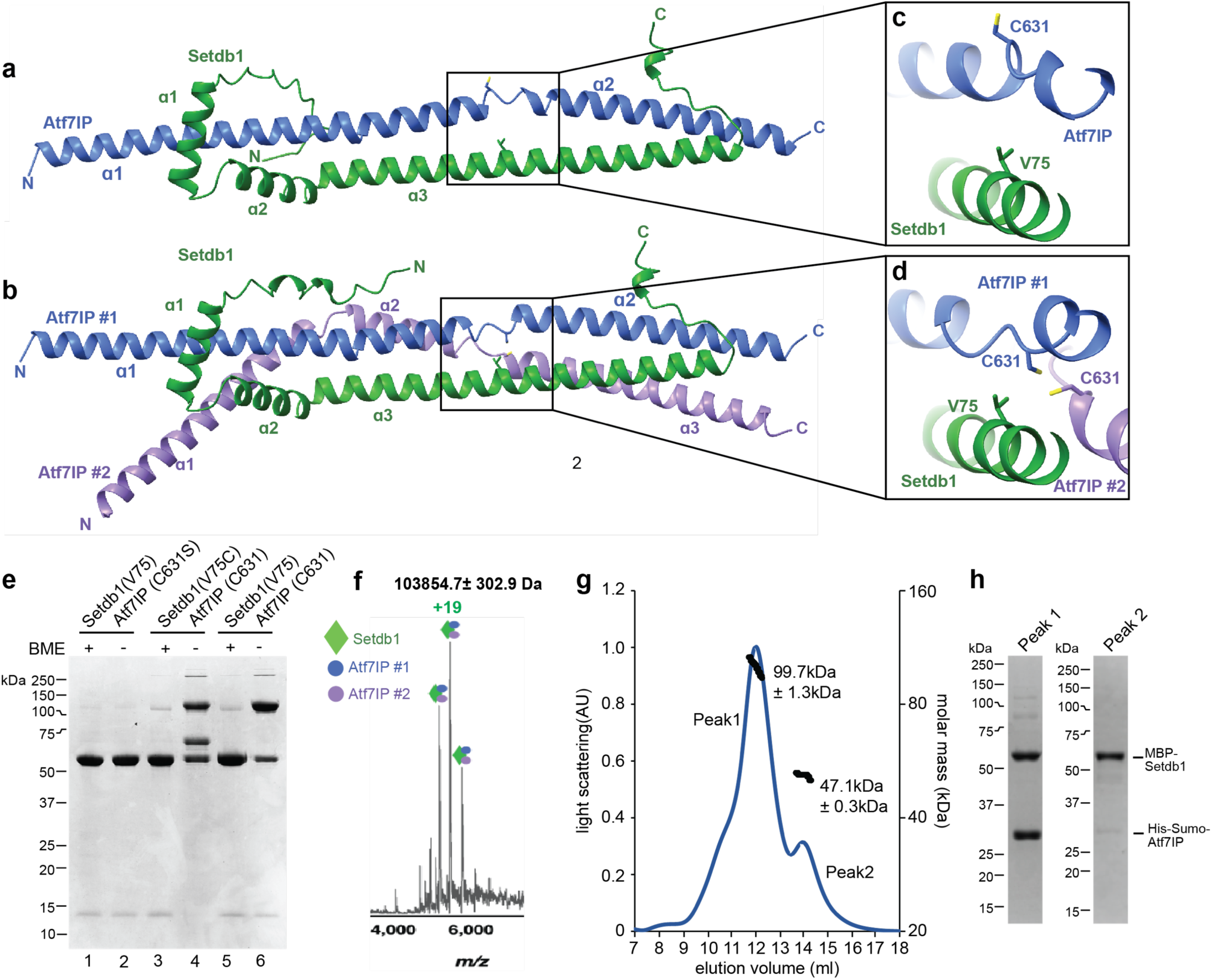
Two molecules of Atf7IP bind to a single Setdb1. a) Alphafold2 prediction of the 1:1 Setdb1:Atf7IP complex. b) Alphafold2 prediction of the 1:2 Setbd1:Atf7IP complex. c-d) close up views of Atf7IP C631 and Setdb1 in the 1:1 and 1:2 Alphafold2 predictions. e) SDS-PAGE gel showing the results of cysteine crosslinking experiments with Setdb1 and Atf7IP Cys mutants. f) native mass spectrum of the intact MBP-Setdb1(2-115)/6Xhis-sumo-Atf7IP(574-666) complex. g) SEC-MALS trace of the intact MBP-Setdb1(2- 115)/6XHis-Sumo-Atf7IP(574-666) complex. h) SDS-PAGE analysis of fractions within peak 1 and peak 2 from the SEC-MALS experiment shown in g.

To validate the Alphafold2 multimer predictions and to determine the stoichiometry of the Setdb1/Atf7IP complex we performed site-specific disulfide crosslinking using a shorter Setdb1(10-108)/Atf7IP(574-666) complex trimmed to include only the residues of Setdb1 with high PLDDT scores (Fig. S2c-d). We utilized the SSbondPre software^32^ to identify optimal sites for engineered Cys mutations predicted to form disulfide crosslinks between Setdb1 and Atf7IP. SSbondPre identified the Setdb1 V75C mutation as a top candidate to form crosslinks with Atf7IP C631. Setdb1 V75 is positioned very close to both copies of Atf7IP C631 in the 1:2 model, but Atf7IP C631 is flipped away from Setdb1 V75 in the 1:1 model (**Fig. 2c,d**). Therefore, we predicted that a Setdb1 V75C mutation would form a disulfide crosslink with Atf7IP C631 in the 1:2 complex, but not in the 1:1 complex. However, we note that Alphafold2 is less confident in the position of Atf7IP C631 in the 1:1 complex, as evident from the lower PLDDT score in this region compared to the rest of the predicted structure (**Fig. S2a,b**). In addition, for the prediction of the 1:2 complex Atf7IP#1 C631 is positioned very close to the Atf7IP#2 C631, suggesting that these residues could form disulfide crosslinks as well, but only in the context of the 1:2 Setdb1/Atf7IP complex (**Fig. 2d**). Therefore, we reasoned that a Setdb1 V75C mutation paired with the native Atf7IP C631 residue could be used to distinguish between the 1:1 and 1:2 complexes predicted by Alphafold2 and validate the position of these residues in the Alphafold2 model.

To accomplish this, we generated three different his-Setdb1(10-108)/MBP-Atf7IP(574-666) complexes containing Setdb1(V75)/Atf7IP(C631S), Setdb1(V75C)/Atf7IP(C631) or Setdb1(V75)/Atf7IP(C631) in a mutant background where all other native cysteine residues were replaced with serine. After purification using the 6xhis affinity tag on Setdb1 in non-reducing conditions we assessed disulfide formation by gel shift in SDS-PAGE (**Fig. 2e**). The Setdb1(V75)/Atf7IP(C631S) complex, which contains no cysteine residues, did not form any apparent disulfide crosslinks and the visible bands corresponded to un-crosslinked his-Setdb1 and MBP-Atf7IP proteins (**Fig. 2e lanes 1&2**). However, for the Setdb1(V75)/Atf7IP(C631) complex, a large band appeared at ∼110kDa which coincided with a reduction of MBP-Atf7IP (**Fig 2e, lane 6**). When this sample was treated with β- mercaptoethanol (BME) we observed a nearly complete depletion of the 110kDa band that was accompanied by a corresponding increase of the MBP-Atf7IP band (**Fig. 2e lane 5**). This behavior is consistent with the 110kDa band being a disulfide linked MBP-Atf7IP(574-666) and suggests that the Setdb1/Atf7IP complex contains at least two copies of Atf7IP, agreeing with the Alphafold2 prediction of the 1:2 complex (**Fig. 2b**). The Setdb1(V75C)/Atf7IP(C631) complex contains the same disulfide-linked MBP-Atf7IP dimer band at 110kDa, suggesting that the Setdb1(V75C) mutation does not alter the underlying stoichiometry of the complex (**Fig. 2e lane 4**). However, for this complex, a new band appeared at 65kDa which coincided with the disappearance of the 6Xhis-Setdb1 band. The 65kDa molecular weight of this band matches the combined molecular weights of 6Xhis-Setdb1(10-108, 13kDa) and MBP-Atf7IP(574-666, 52kDa) and when this sample is treated with BME, the 110kDa and 65kDa bands are depleted, while the intensity of MBP-Atf7IP and 6Xhis-Setdb1 bands are increased (**Fig. 2e lane 3**). This is consistent with the 65kDa band being a disulfide linked 6Xhis-Setdb1(10-108)/MBP- Atf7IP(574-666) dimer and supports the Alphafold2 model which places Setdb1 V75 close to Atf7IP C631 in the 1:2 complex.

To further validate the stoichiometry of the Setdb1/Atf7IP complex, we performed native mass spectrometry on a MBP-Setdb1(2-115)/6Xhis-SUMO(HS)-Atf7IP(574-666) complex. Mass spectra of the intact complex showed a species corresponding to mass 103854.7 ± 302.9 Da (**Fig. 2f, S4, left panel**) which matches the predicted mass of a 1:2 MBP-Setdb1(2-115):HS-Atf7IP(574-666) complex at 102,620.23 Da (with the extra experimental mass corresponding to adducting of salts and solvent, as is typical with native MS when avoiding complex dissociation and restructuring). Additionally, when higher-energy collision dissociation (HCD) fragmentation was carried out by isolating a 500 m/z (m/z 5300 to 5800) window with quadrupole, and subsequent application of HCD 220 V, two fragments were observed at HCD 220 (**Fig S4, right panel**). In the high m/z range, a fragment with mass 80327.0 ± 193.1 Da was observed and this species likely contains one copy of Setdb1 and one copy of Atf7IP (Expected mass = 79658.09 Da). In the low m/z range, a smaller species with mass 23255.3 ± 21.9 Da was observed, corresponding to one copy of Atf7IP. As collision induced dissociation fragments a complex along the weakest interface, it is possible that the Setdb1:Atf7IP complex has a higher binding affinity, while the second copy of Atf7IP is bound more weakly (**Fig S4, left panel**).

Finally, analysis of the same complex containing MBP-Setdb1(2-115, 56.7 kDa) and HS-Atf7IP(574-666, 23.2 kDa) using Size Exclusion Chromatography with Multi Angle Light Scattering (SEC-MALS) showed that the MBP- Setdb1(2-115)/HS-Atf7IP(574-666) complex has a molecular weight of ∼100kDa (**Fig. 2g**), which agrees with the expected molecular weight of a 1:2 complex between these proteins (103 kDa). A second peak with a molecular weight of ∼47 kDa, corresponds to isolated MBP-Setdb1 that was co-purified with the complex following MBP affinity purification (**Fig 2h**). Together, these data clearly indicate that Setdb1 and Atf7IP form a heterotrimer with a 1:2 Setdb1:Atf7IP stoichiometry, agreeing with the Alphafold2 prediction of the 1:2 complex.

### The Setdb1/Atf7IP complex is multimeric in cells

To rule out that the minimal Setdb1/Atf7IP complex forms a 1:2 heterotrimer due to the truncations used in the biochemical experiments described above or due to heterologous expression in *E. coli*, we investigated the stoichiometry of the full-length Setdb1/Af7IP complex expressed in HEK 293 cells. We co-expressed combinations of FLAG-Atf7IP, Myc-Atf7IP and HA-Setdb1 and immunoprecipitated the Setdb1/Atf7IP complexes using Anti-FLAG magnetic beads. We reasoned that if Setdb1 and Atf7IP form a 1:2 complex in cells, Myc-Atf7IP should co-IP with FLAG-Atf7IP. However, if Setdb1:Atf7IP form a 1:1 complex, no Myc-Atf7IP should co-IP with FLAG-Atf7IP (**Fig 3a**). When FLAG-Atf7IP, Myc-Atf7IP and HA-Setdb1 were co-expressed, HA-Setdb1 and Myc- Atf7IP pulled down with FLAG-Atf7IP (**Fig 3b, lane1**). The co-IP of Myc-Atf7IP suggests the full length Setdb1/Atf7IP complex contains more than one copy of Atf7IP, clearly agreeing with the 1:2 binding stoichiometry observed for the minimal Setdb1/Atf7IP complex (**Fig. 2**). Importantly, HA-Setdb1 and Myc-Atf7IP did not pulldown with FLAG-Atf7IP(Δ574-666), which lacks the coiled coil region that binds to Setdb1 (**Fig 3b, lane2**), or from cells with Atf7IP lacking the FLAG affinity tag (**Fig. 3b, lane 4**). Surprisingly, Myc-Atf7IP pulled down with FLAG-Atf7IP even in the absence of co-expressed HA-Setdb1 (**Fig. 3b, lane 4**). The association between FLAG- Atf7IP and Myc-Atf7IP was not due to co-IP of endogenous untagged Setdb1, as no endogenous Setdb1 was observed in the FLAG pulldown (**Fig. 3b, lane 4**). These results indicate that the native Setdb1/Atf7IP complex contains at least two copies of Atf7IP and that Atf7IP can form a multimeric complex with itself.

**Figure 3:**
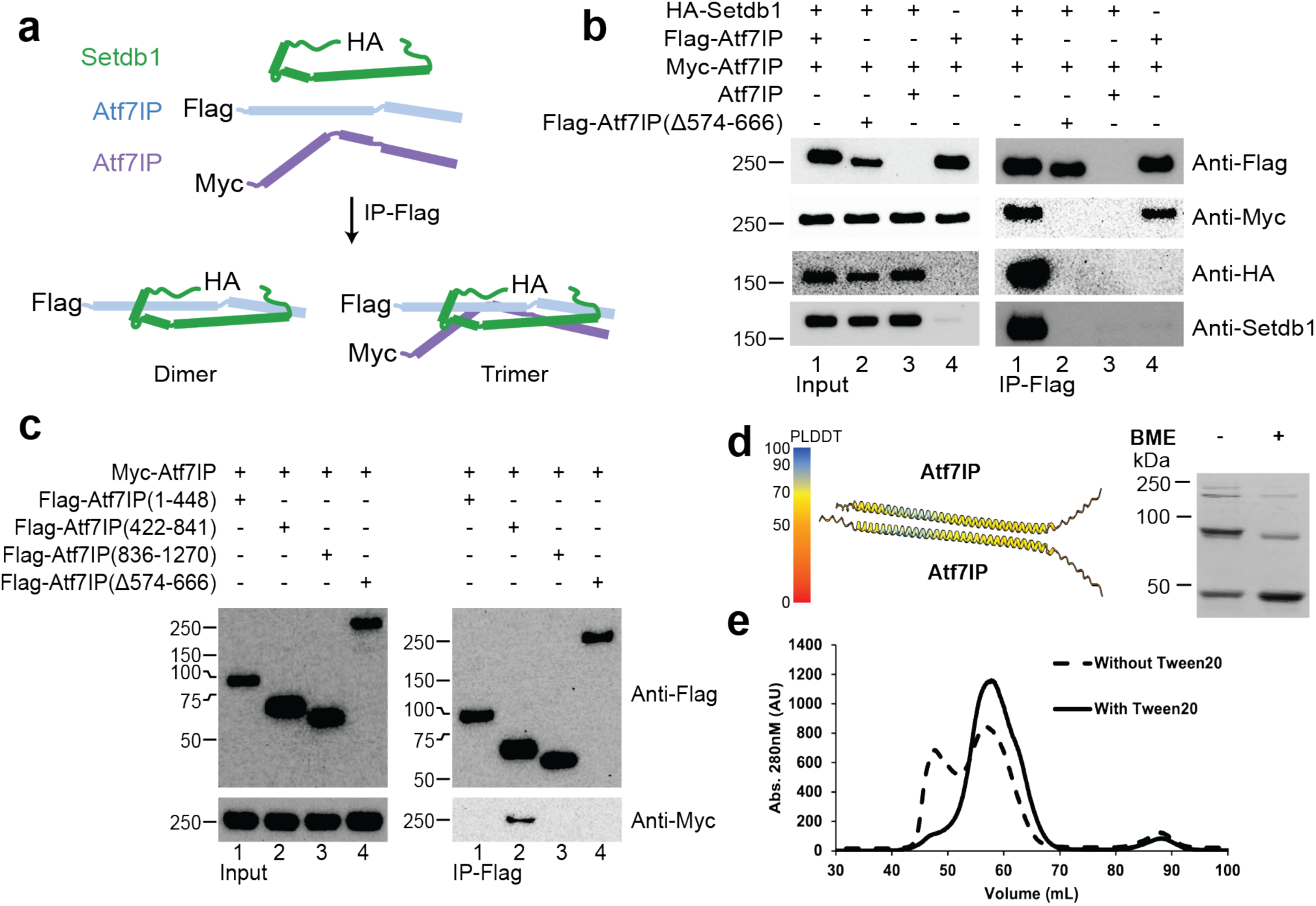
Setdb1/AF7IP form a mulKmeric complex in cells. a) Pulldown design that allows discriminaKon between dimer and mulKmer Setdb1/Am7IP complexes. b-c) FLAG-pulldowns from HEK293 cells expressing the indicated proteins. d) CharacterisKc Alphafold2 predicKon of the Am7IP(574-666)/Am7IP(574-666) complex showing poor accuracy (le|), cysteine-cysteine crosslinking assay of mutant MBP-Am7IP (574-666) C608S C612S with and without Beta-mercaptoethanol (BME). e) Gel FiltraKon assay of MBP-Am7IP (574-666) with and without Tween20.

To further investigate the self-association of Atf7IP, we co-expressed FLAG-tagged truncations of Atf7IP with full length Myc-Atf7IP followed by immunoprecipitation using the FLAG tag. Only Atf7IP(422-841), which contains the coiled coil domain was able to co-IP with Myc-Atf7IP (**Fig. 3c, IP-Flag; lane 2**). FLAG-Atf7IP(Δ574-666) did not pulldown with Myc-Atf7IP, suggesting that the coiled coil region is essential for Atf7IP self-association in cells. To understand how the 574-666 region of Atf7IP self-associates, we generated models of Atf7IP(544-677) homomeric complexes of different stoichiometries using Alphafold2 multimer (**Fig 3d; left panel, S3**). However, Alphafold2 did not predict any multimeric structure of Atf7IP(544-677) with high confidence suggesting that the homomeric Atf7IP complex may have a heterogeneous stoichiometry prior to its interaction with Setdb1.

This is supported by disulfide crosslinking experiments using a cysteine-free mutant of MBP-Atf7IP(574-666) containing only the native C631. In Isolation, MBP-Atf7IP(574-666, C631) formed multiple high molecular weight bands on the SDS page under non-reducing conditions. These high molecular weight bands were partially resolved upon treatment with BME. This suggests that MBP-Atf7IP(574-666) can form SDS-resistant high molecular weight complexes. (**Fig. 3d; right panel**). In addition, size exclusion profiles of the isolated Mbp- Atf7IP(574-666) fragment show a significant portion of the protein shifted toward the void volume (**Fig. 3e**), indicating that isolated Atf7IP(574-666) can generate larger multimeric complexes. These high molecular weight multimers of Atf7IP can be resolved by including Tween 20 in the size exclusion buffer (**Fig. 3e**), or by co- expression with Setdb1 (**Fig. 2g**) indicating that Atf7IP homomeric complexes may non-specifically self-associate until bound by Setdb1. Together these data indicate that Atf7IP forms multimeric complexes *in vitro* and can self- associate in cells.

### Atf7IP directly competes with CRM1 for Setdb1 binding

A key function of Atf7IP is to prevent removal of Setdb1 from the nucleus by the CRM1 nuclear export machinery^18^. The N-terminal region of Setdb1 contains two putative class 1c NES motifs^33^ that are thought to be responsible for its nuclear export (**Fig 1a**)^25, 34^. In the 1:2 Setdb1:Atf7IP Alphafold2 model, both Setdb1 NES motifs directly interact with Atf7IP#1 and Atf7IP#2 (**Fig 4a**). NES #1 spans residues 23-36 and is located within α1 and the loop between α1 and α2. NES #2 spans residues 99-105 and is located at the C-terminal end of α3. Hydrophobic residues of both Setdb1 NES motifs contact each copy of Atf7IP and are predicted to form multiple hydrophobic interactions. NES#1 is predicted to contact both copies of Atf7IP near residues 591-602 (**Fig. 4b**) while NES #2 contacts each copy of Atf7IP near residues 649-660 (**Fig. 4c**). The extensive hydrophobic interactions mediated by the Setdb1 NES motifs may contribute to high affinity binding between Setdb1 and Atf7IP. This agrees with recent reports that the NES motifs of Setdb1 are required for the interaction between Setdb1 and Atf7IP in cells^25^. Deletion of Setdb1 α1, which encompasses NES#1, moderately reduced binding between Setdb1 and Atf7IP to ∼80% of WT as measured by Alphascreen (**Fig. 4d**). However, a larger deletion of Setdb1 α1 and α2 reduced Atf7IP binding to ∼30% of WT. This indicates that while NES#1 contributes to Atf7IP binding, tight binding between Atf7IP and Setdb1 depends more on the overall size of the helical interaction between Setdb1 and Atf7IP than on the specific interactions with the NES motifs.

**Figure 4:**
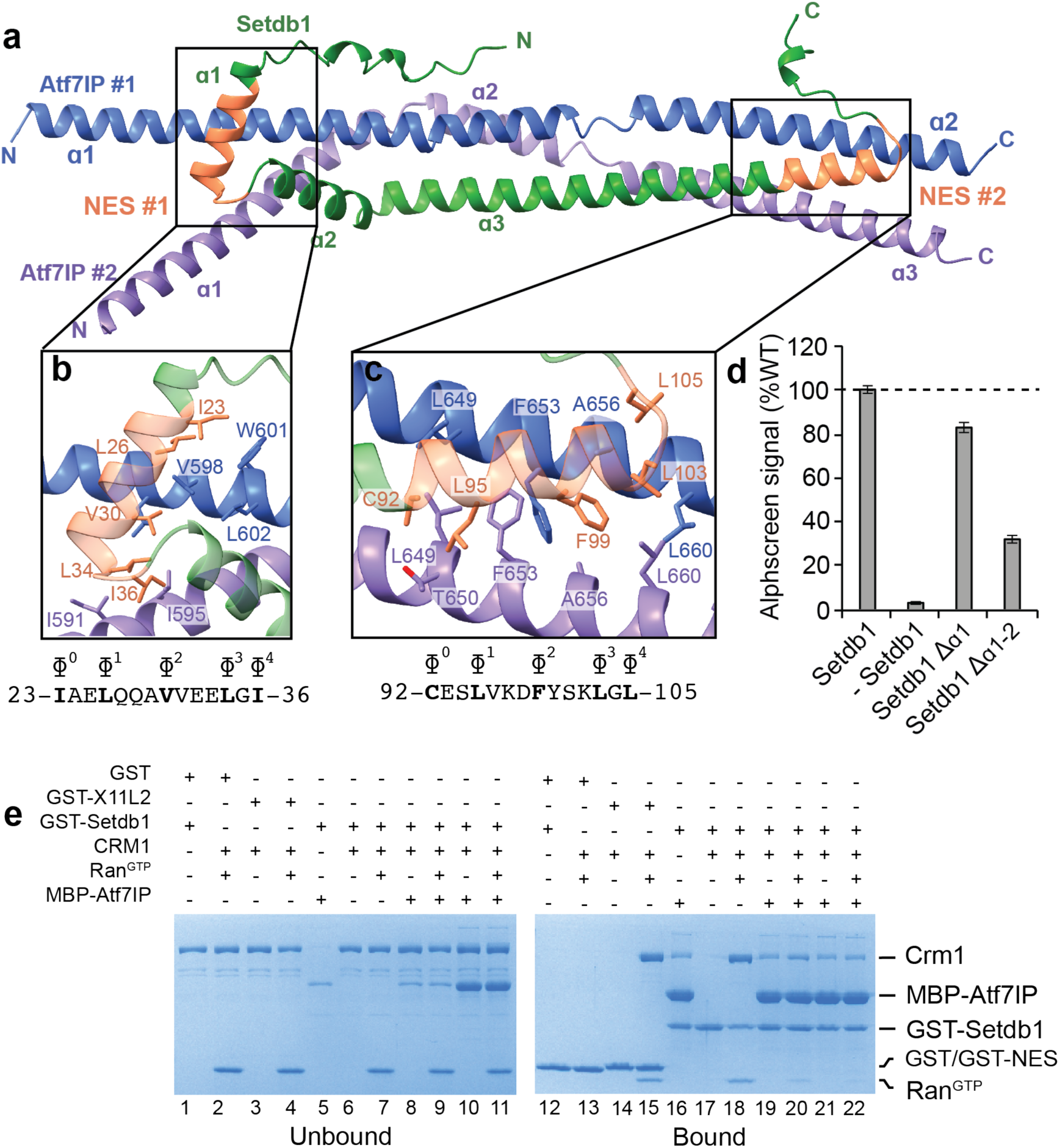
ATF7IP binds to Setdb1 NES motifs and blocks CRM1 binding. **a**) Alphafold prediction of the 1:2 Setdb1:Atf7IP heterotrimer. The location of the Setdb1 NES motifs are indicated and colored orange. **b-c**) Top: close up views of the Setdb1 NES motifs showing detailed interactions between each NES and each copy of Atf7IP. Critical NES residues of Setdb1 and interacting residues of Atf7IP are shown as sticks. Bottom: Sequence of each Setdb1 NES motif with the critical hydrophobic NES residues indicated with ɸ. **d**) Alphascreen binding assay between MBP-Atf7IP(574-666) and the indicated his-sumo-Setdb1(2-114) with the indicates truncations. **e**) GST pulldown assays showing competition for Setdb1 binding between Atf7IP and Crm1.

The direct interactions between the Setdb1 NES motifs and Atf7IP predicted by Alphafold2 suggest that Atf7IP inhibits nuclear export of Setdb1 by blocking CRM1 binding. To determine if Atf7IP can directly compete with CRM1, we assessed binding between GST-Setdb1(2-115) and CRM1 in the absence or presence of MBP- Atf7IP(574-666). CRM1 pulled down with GST fused to the known X11L2 NES motif (GST-X11L2) and with GST- Setdb1 (2-115) immobilized on glutathione beads in a Ran^GTP^ dependent manner (**Fig. 4e, lane 3-4, 6-7**, **14-15, 17-18**), validating that the N-terminus of Setdb1 has *bone fide* NES motifs that CRM1 can recognize *in vitro*. In agreement with our prior binding assays (**Fig. 1**), MBP-Atf7IP (574-666) also strongly pulled down with GST- Setdb1 (2-115) (**Fig. 4e, lane 16**). However, pre-incubation of GST-Setdb1(2-115) with a 3-fold excess of MBP- Atf7IP(574-666) reduced binding of CRM1 in the presence of Ran^GTP^ (**Fig. 4e lane 20**). Similar results were observed when 15-fold molar excess of MBP-Atf7IP(574-666) was used, which completely blocked CRM1 from binding to Setdb1. (**Fig 4e lane 22**). Therefore Atf7IP(574-666) directly competes with CRM1 for binding to the NES motifs within Setdb1(2-115), providing a structural rationale for Atf7IP-dependent nuclear retention of Setdb1.

### Atf7IP and Atf7IP2 use conserved structural features to bind Setdb1

Recent studies have shown that Atf7IP is replaced by its paralog Atf7IP2 in mouse germline cells and is required for spermatogenesis^27, 35^. In these cells Atf7IP2 functions analogously to Atf7IP by localizing Setdb1 to the nucleus, blocking its nuclear export and recruiting Setdb1 to specific chromatin loci for deposition of H3K9me2-3^27^. Like Atf7IP, the middle region of Atf7IP2, spanning residues 328-380, is predicted to have strong coiled coil character^28^ (**Fig. 5a**). Atf7IP and Atf7IP2 share a sequence identity of 15% (**Fig. S5**), while the coiled coil regions share an identity of 24% (**Fig. S5, 5b**). Because coiled coils are defined by a simple pattern of hydrophobic residues, homologous coiled coil proteins can have highly degenerate sequences^36^. Indeed, even though the coiled coils of Atf7IP and Atf7IP2 have low sequence identity, several interfacial hydrophobic residues are conserved, while others are substituted with similar amino acids (**Fig 5b**). We therefore asked if the Atf7IP2 coiled coil region and the N-terminus of Setdb1 adopt a similar structure to the Atf7IP/Setdb1 complex described above. We used Alphafold2 to predict structures of Setdb1(2-115) and Atf7IP2(297-388) in 1:1 and 1:2 Setdb1:Aft7IP2 stoichiometries (**Fig. 5c-d, S6a-b**). Both structures of Setdb1/Atf7IP2 were predicted with high confidence and are superimposable with predictions of Sedb1/Atf7IP, aligning with RMSDs of 0.95Å for the 1:1 structures and of 1.04Å for the 1:2 structures (**Fig. S6c-d**). Since Alphafold2 predicted 1:1 and 1:2 Setdb1/Atf7IP2 complexes with similar confidence we wanted to determine if the 1:2 stoichiometry of the Setdb1/Atf7IP complex is also conserved in the Setdb1/Atf7IP2 complex. Therefore, we co-expressed MBP- Setdb1(2-115), HS-Atf7IP2(297-388) and Strep-Atf7IP2(297-388) in *E. coli* and purified the resulting complex sequentially using the His and Strep tags on both copies of Atf7IP2 (**Fig 5e**). After the sequential affinity purification all three proteins eluted together, indicating that Setdb1 and Atf7IP2 form a multimeric complex in agreement with the 1:2 Setdb1:Atf7IP2 model that was predicted by Alphafold2(**Fig. 5d, f**). Therefore, Atf7IP2 binds to Setdb1 to in a 2:1 stoichiometry using a conserved interface that is shared with Atf7IP Because Alphafold2 predictions of the minimal Setdb1/Atf7IP2 complex closely matched predictions of the Setdb1/Atf7IP complex, we wondered if Atf7IP and Atf7IP2 could form mixed heterotrimeric complexes with single copies of Setdb1, Atf7IP and Atf7IP2. Alphafold2 predictions of the Setdb1/Atf7IP/Atf7IP2 heterotrimeric complex showed a similar arrangement of Atf7IP and Atf7IP2 as predicted for the Setdb1/Atf7IP and Setdb1/Atf7IP2 complexes (**Fig. 5g**). Interestingly, in the Setdb1/Atf7IP/Atf7IP2 heterotrimeric complex Alphafold2 always placed Atf7IP and Atf7IP2 in the same position, potentially suggesting that the two proteins are only compatible in a single orientation with Setdb1 (**Fig. S6c**). Therefore, we co-expressed MBP-Setdb1(2- 115), HS-Atf7IP(574-666) and Strep-Atf7IP2(297-388) in *E. coli* and sequentially purified the complex using the his and strep affinity tags on Atf7IP and Atf7IP2 (**Fig. 5h**). In agreement with the Alphafold2 Setdb1/Atf7IP/Atf7IP2 model, the sequential purification yielded complexes that contained HS-Atf7IP2, Strep-Atf7IP2 and MBP-Setdb1 (**Fig. 5i**) indicating that Setdb1 can readily form mixed complexes with Atf7IP2 and Atf7IP. This indicates that the precise composition of Setdb1/Atf7IP/Atf7IP2 complexes may have important roles in the targeting and regulation of Setdb1.

**Figure 5:**
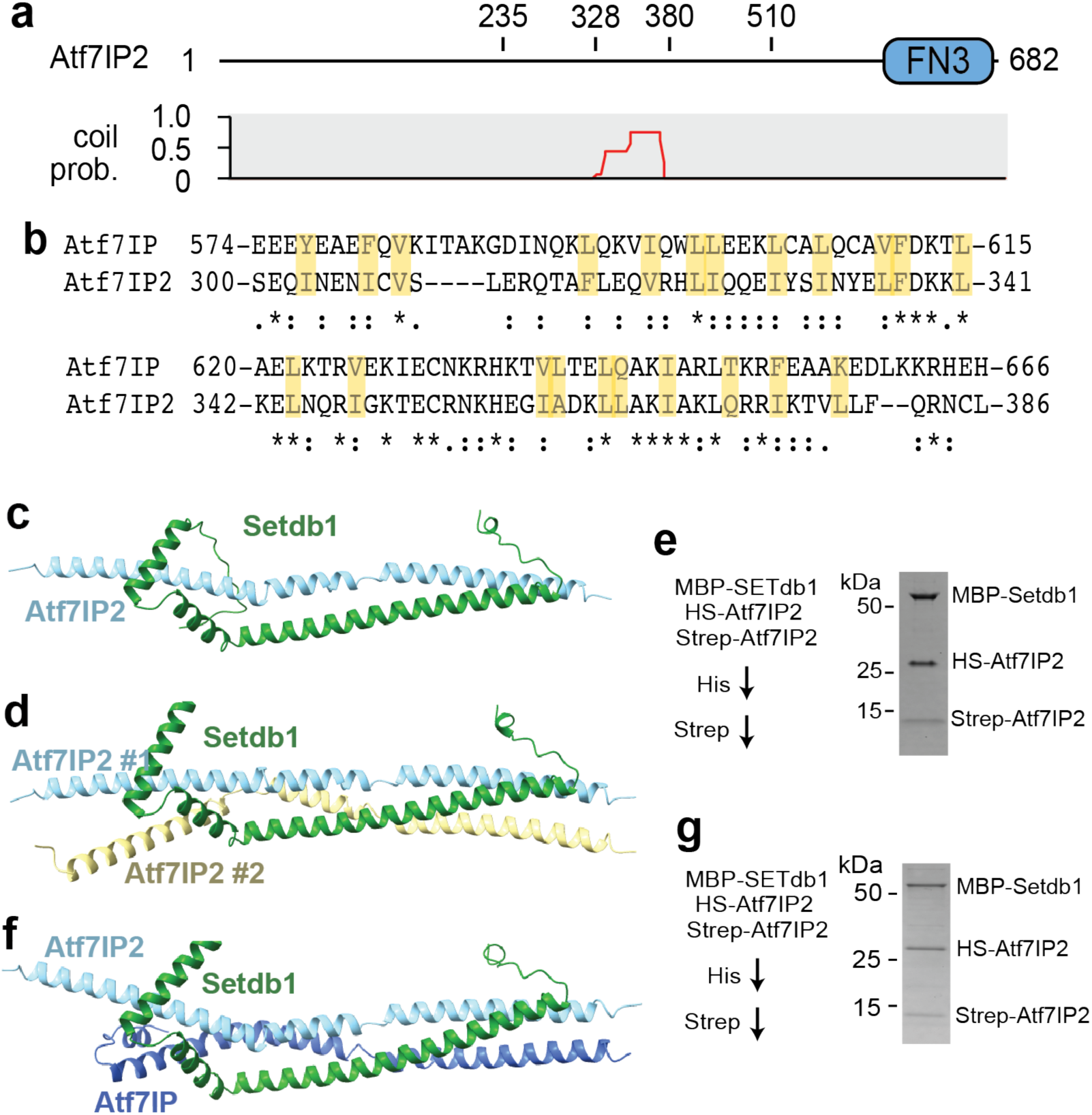
tf7IP2 mimics Atf7IP heterotrimeric interaction with SETdb1. a) Domain architecture of Atf7IP2 (top) with the corresponding pcoils prediction (bottom). FN3 = fibronectin type 3 domain. b) Sequence alignment showing the similarity of coiled-coiled regions of Atf7IP and Atf7IP2. c-d) Alphafold2 predictions of the 1:1 Setdb1(2-114):Atf7IP2(297- 388) complex and the 1:2 Setdb1(2-114):Atf7IP2(297-388) complexes e) Sequential His and Strep pulldown design that allows the discrimination of the MBP-Setdb1(2-114)/HS- Atf7IP2(297-388) dimer from the MBP-Setdb1(2-114)/HS-Atf7IP2(297-388)/Strep-Atf7IP(297- 388) heterotrimer. f) SDS-PAGE gel results of experiments in e) showing formation of heterotrimer. g) Alphafold2 predictions of the mixed trimer; Setdb1(2-114)/Atf7IP(574- 666)/Atf7IP2(297-388) complexes. h) Sequential His and Strep pulldown design that allows the discrimination of the MBP-Setdb1(2-114)/HS-Atf7IP(574-666) dimer from the MBP-Setdb1(2- 114)/HS-Atf7IP(574-666)/Strep-Atf7IP2(297-388) mixed heterotrimer i) SDS-PAGE gel results of experiments in h) showing formation of mixed trimer. HS; 6Xhis-SUMO.

## Discussion

Our study provides the first structural and biochemical evidence showing that the N-terminal helical region of Setdb1 directly binds to the central helical fragment of Atf7IP by forming a highly intertwined three-helix coiled- coil. The Alphafold2 models of the complex between Atf7IP and Setdb1 show that critical nuclear export signals within Setdb1 are completely blocked by Atf7IP, preventing CRM1 binding to Setdb1. This explains several prior observations showing that Setdb1 nuclear localization is completely dependent on ATF7IP and that its nuclear export relies on the CRM1 export machinery^16, 33, 37, 38^. Therefore, our study provides a structural rationale to explain the cellular localization of Setdb1.

Setdb1 over-expression in cancers enhances immune evasion by restricting antigen presentation of retrotransposon-derived peptides^39^. Because the interaction with Atf7IP is critical for Setdb1-mediated silencing of retrotransposons, disrupting this interaction may be an attractive strategy to reduce immune evasion of cancers^40^. Setdb1 and Atf7IP interact through coiled-coils and targeting this 1:2 complex with peptides or compounds that block Setdb1/Atf7IP binding could be open new areas for cancer treatment [cit]. The finding that Setdb1 and Atf7IP interact with a 1:2 Setdb1:Atf7IP stoichiometry was unexpected because in most human tissues the expression of Setdb1 and Atf7IP closely matched (**Fig. S7**). However, this binding stoichiometry could explain several observations showing that a significant fraction of Setdb1 is retained in the cytosol^18, 41, 42^. Indeed, Setdb1 has several is cytosolic substrates and binding partners that are important for normal cellular processes and development of cancer^43–45^. If Atf7IP and Setdb1 formed an exclusive 1:1 complex this would mean that most Setdb1 would be bound to Atf7IP and localized to the nucleus. However, the 1:2 Setdb1:Atf7IP binding stoichiometry suggests that complete nuclear localization of Setdb1 would require twice as much Atf7IP as Setdb1. Therefore, we propose that the 1:2 binding stoichiometry of Setdb1:Atf7IP allows the cell to control the amount of Setdb1 in the cytosolic and nuclear fractions, ensuring that a fraction of Setd1 remains cytosolic.

Setdb1’s ability to form 1:2 heterotrimers with Atf7IP or Atf7IP, and its ability to form mixed 1:1:1 heterotrimers with Atf7IP and Atf7IP2 suggests that formation of these different complexes may serve specific functions in transcription regulation. In the testis Atf7IP2 directs Setdb1 to testis-specific chromatin loci that regulate meiosis and sex chromosome inactivation^27, 35^. In addition, Atf7IP recruits Setdb1 to various co-repressor complexes that silence retrotransposons [MBD1, Trim28, HUSH, Mettl1, KDM5b]. Atf7IP and Atf7IP2 are expressed together in many tissues, indicating that that mixed 1:1:1 timers between Setdb:Atf7IP:Atf7IP2 could potentially form in many contexts (**Fig. S7**). Indeed, Atf7IP and Setdb1 localization can be disrupted in testis following knockout of Atf7IP2^35^, indicating that there is some crosstalk between these two Atf7IP paralogs. Therefore, the localization and activity of Setdb1 may be fine-tuned by formation of tissue-specific Atf7IP/Atf7IP2/Setdb1 complexes, potentially contributing to the tissue specific functions of Setdb1 in rTE silencing.

## Author Contributions

E.J.W. and L.K. conceived of the study. L.K., K.P. and A.B.N. conducted the biochemistry experiments. I.M. did the mass-spectrometry experiments. L.K. and E.J.W. wrote the manuscript. E.J.W., Y.M.C. and V.H.W. supervised the study.

## Materials and Methods Cloning and protein expression

Human Setdb1 and Atf7IP were PCR amplified from cDNAs #OHu26362 (genscript) and ORFeome_LaBaer- 011.0029 (DNASU) and cloned into the pLib donor vector from the BigBac insect cell expression system^46^. We also used the genescript New MB Gene Fragments service to generate the Strep-Atf7IP2(297-388) DNA fragment with overhangs needed for cloning. Fragments of Setdb1, Atf7IP and Atf7IP2 were individually cloned into pETduet-1 or pET28a or pACYC vectors containing N-terminal 6xHis, 6xHis-Sumo, AviTag-Mbp, GST or strep affinity tags (table S1). Co-expression vectors were constructed by cloning tagged fragments of Setdb1 and Atf7IP or Setdb1 and Atf7IP2 into the first and second multiple cloning sites of pETduet-1.

For cysteine-cysteine crosslinking, we used the Genscript gene synthesis and New MB Gene fragments services to generate a pETduet-1 co-expression vector containing 6xHis-Setdb1 C53S C92S (10-108)/ATF71P C608S C612S (574-666) where all native cysteine residues were mutated to serine except Atf7IP(C631) and the gene fragment At7IP2 C308S C385S (297-388). All mutants of Atf7IP(C631), Setdb1(V75) and Atf7IP2(C353) were generated from the aforementioned vector and gene fragment using site directed mutagenesis.

Full-length Setdb1 and Atf7IP and various truncations of Atf7IP were also individually cloned into pEG BacMam vectors containing N-terminal FLAG, Myc and HA tags (table S1) for transient expression in HEK293 cells (described in more detail in the method section on the in vivo pull-down and immunoprecipitation assays).

GST and GST-X11L2^NES^ were expressed in pGexTev vector in BI21 gold cells and purified according to established protocols^37^. The proteins hCRM1 and Ran^GTP^ were also expressed and purified according to previously published protocols^47^.

All other vectors were individually transformed or co-transformed with pEWL17 (containing Strep-Atf7IP2(297- 388)) into BL21DE3 or Rosetta2 cells and grown in LB media containing 100 µg/mL ampicillin and/or 25µg/mL of chloramphenicol and/or 100µg/mL kanamycin (depending on the antibiotic resistance gene in the plasmids used for transformation) at 37°C until the cell density reaches an OD_600_ of 0.6–0.8. At this point, protein expression in large scale cultures (2L) was induced with 1 mM IPTG, and cells were incubated for 16 hours at 18°C. The cultures expressing Avi-tag MBP-Atf7IP(574-666)/birA were additionally supplemented with 40µM biotin to express the biotinylated Avi-tag-MBP-Atf7IP(574-666) needed for the alpha screen experiments. When proteins were expressed in small-scale cultures(15-50ml), the Novagen Overnight Express™ Autoinduction System was used.

All cultures were harvested by centrifugation. The pellets of most large cultures (2L) were resuspended in 25ml of lysis buffer containing 30mM HEPES pH 7.9, 400mM NaCl, 10% glycerol and 0.05% Tween-20 and 2mM BME and stored at -80°C until ready to purify. The pellets of small cultures were flash-frozen without resuspension in lysis buffer and stored at -20°C for immediate use the following day. The pellets of cultures that expressed proteins for alpha screen assays were resuspended in 25mL lysis buffer containing 25mM HEPES pH 8, 200mM NaCl and 5mM Beta-mercaptoethanol (BME) prior to storage at -80°C.

## Protein purification

All frozen cell pellets were thawed at 25°C and cooled on ice. 100µM phenylmethylsulfonyl fluoride (PMSF), 20- 25mM Imidazole (for His-tagged proteins) and one protease inhibitor cocktail tablet (Sigma-Aldrich) was stirred into 50-100mL of resuspended pellet in lysis buffer. All cells were lysed by a laboratory homogenizer (SPXFLOW) or by sonication.

The clarified lysates from the cells that expressed 6Xhis-SUMO-Setdb1(2-114), 6Xhis-SUMO-Setdb1(Δ⍺1; residues 35-114), 6Xhis-SUMO-Setdb1(Δ⍺1- ⍺2; residues 50-114) were each loaded onto a 5mL Cytiva HisTrap HP prepacked column pre-equilibrated with 25mM HEPES pH 8, 200mM NaCl and 5mM BME. The loaded column was washed with 50mL of lysis buffer and the proteins were eluted with lysis buffer supplemented with 500mM Imidazole. The eluted proteins were then further purified by dialysis against buffer containing 20mM HEPES pH8 and 200mM NaCl at 4°C overnight.

The cell pellets from cultures that expressed biotinylated Avi-tag-MBP-Atf7IP(574-666), GST-Setdb1(2-114), 6Xhis-Setdb1(11-104)/MBP-Atf7IP(585-666) and MBP-Setdb1(2-115)/6Xhis-sumo-Atf7IP(574-666) were resuspended in 30mM HEPES pH 7.9, 400mM NaCl, 10% glycerol, 0.05% Tween-20, 2mM BME, 1 mM PMSF, and protease inhibitor cocktail, lysed and clarified as described above.

The clarified lysate containing biotinylated Avi-tag-MBP-Atf7IP(574-66) and MBP-Setdb1(2-115)/6Xhis-sumo- Atf7IP(574-666) were loaded onto 5mL (Cytiva) MBPTrap HP prepacked columns pre-equilibrated with lysis buffer and then eluted with lysis buffer supplemented with 10mM Maltose.

The clarified lysate containing 6Xhis-Setdb1(11-104)/MBP-Atf7IP(585-666) was loaded onto a 5mL Cytiva HisTrap HP prepacked column pre-equilibrated with lysis buffer supplemented with 500mM Imidazole.

The clarified lysate containing GST-Setdb1(2-115) were loaded onto 5mL (Cytiva) GSTTrap HP prepacked columns pre-equilibrated with lysis buffer and then eluted with lysis buffer supplemented with 10mM Glutathione.

These proteins were further purified using size-exclusion chromatography on a superdex200 column with 30mM HEPES pH 7.9, 400mM NaCl, 10% glycerol, 0.05% Tween-20, 2mM BME, except GST-Setdb1 which was purified with 20mM HEPES, 100mM KOAc, 2mM MgOAc,10% glycerol and 2mM Dithiothreitol (DTT). The purity of all proteins was assessed by SDS-PAGE, and protein concentrations were determined using a nanodrop. The proteins were spin-concentrated, aliquoted, flash-frozen in liquid nitrogen and stored at -80°C.

## AlphaFold2 Prediction of Setdb1 and ATF7IP Interaction

We used Alphafold2 (AF2 v2.3.1)’s multimer functionality, which incorporates both intra- and inter-molecular constraints to predict model interactions between Setdb1 and ATF7IP. The protein sequences used as input were human Setdb1 (UniProt: Q15047) and ATF7IP (UniProt: Q6VMQ6), retrieved from UniProt. The sequences of Setdb1 and ATF7IP were concatenated for complex prediction without a linker, as AF2 handles independent chains. Multiple sequence alignments (MSAs) were generated using the built-in AlphaFold pipeline, pulling data from sequence databases like UniRef and MGnify. Five models were generated, and the highest-confidence model was selected based on the predicted local distance difference test (pLDDT) and predicted aligned error (PAE) scores. A pLDDT score threshold of >70 was used to identify well-resolved regions and assess model confidence in the interaction interface. The predicted complex was analyzed in ChimeraX with particular attention given to potential interaction interfaces. Residue-residue contacts within 5 Å at the interface were identified as potential interaction sites. The Setdb1 and Atf7IP input sequences fed to AF2 were trimmed iteratively to obtain minimal interface of the Setdb1-Atf7IP interaction and multiple stoichiometric combinations were tested to evaluate model confidence.

## Validation assays for AF2 models predicting Setdb1/Atf7IP interaction Gel-Filtration binding assays

Recombinant 6Xhis-SUMO-Setdb1(2-115) and MBP-Atf7IP(574-666) proteins were individually expressed and purified as described above. The purified 6Xhis-Setdb1 at 40µM was mixed with purified Avi-tag-MBP- Atf7IP(574-666) at 20µM in 307.8µl of binding buffer (30mM HEPES pH 7.9, 400mM NaCl, 10% glycerol, 2mM BME and 0.05% Tween-20) and incubated for 1 hour at room temperature to allow binding interactions to occur. Control samples of individual proteins were prepared similarly. The protein mixture and control samples were loaded sequentially onto the same Superose6 column (Cytiva) which was pre-equilibrated with the binding buffer. The column was re-equilibrated after every use and elution of each sample was monitored by absorbance at 280 nm. Fractions of 0.5 mL were collected throughout the run and analyzed by SDS page gel electrophoresis alongside their gel filtration profiles.

## Alphascreen assays

To assess the strength of the interactions between 6Xhis-SUMO-Setdb1(2-115) and biotin-MBP-Atf7IP(574- 666), an Alphascreen^29^ assay was performed using the Alphascreen His-Detection kit (PerkinElmer) according to the manufacturer’s instructions. The purified 6Xhis-SUMO-Setdb1 and AviTag-MBP-Atf7IP were first used at equimolar concentrations of 20nM diluted in 1X Alphascreen assay buffer (50mM MOPS, 50mM CHAPS, 50mM NaF, 100µg/ml BSA, pH7). For following experiments, titrations were also performed keeping one protein at 20nM while varying the concentration of the other protein from 0-20nM. Each reaction was done in triplicates and was incubated for 2 hours at room temperature before adding the beads. After the initial incubation, both Nickel Chelate Alphascreen Acceptor beads (PerkinElmer) and Streptavidin-coated Donor beads (PerkinElmer) were added at 5 µg/mL per bead type to each reaction. Each reaction (40µl) was then incubated in the dark for another hour at room temperature and transferred to a 384-well white OptiPlate (PerkinElmer). Following incubation, the Alphascreen signal was measured using an EnVision Multilabel Plate Reader (PerkinElmer) with excitation at 680 nm and emission detection at 520-620 nm. The signal was recorded as counts per second (CPS), which is proportional to the proximity of the His- and biotin-tagged proteins brought together by interaction.

To quantify the interactions, the raw Alphascreen CPS values for each triplicate was expressed as mean ± standard deviation and plotted against protein concentrations, and binding affinities (apparent dissociation constant, Kd) were calculated by fitting the data to a one-site binding model using GraphPad Prism software (v9.0). Wells containing only His-SUMO-Setdb1(2-114) or Biotin-MBP-Atf7IP(574-666) without their interaction partner were used as negative controls, and the biotinylated- 6Xhis positive control from the Alphascreen His- Detection kit (PerkinElmer) was also used in each experiment.

## Site-specific cysteine crosslinking assays

We used SSbondpre, a computational method based on a neural network (trained with structures curated from the Protein Data Bank) to predict disulfide bond formation between Atf7IP(C631) and Setdb1(V75) and with itself in the Setdb1(2-115)/Atf7IP(574-666) protein complex AF2 model. By further evaluating the AF2 models of the trimer complexes between Setdb1, Atf7IP, and Atf7IP2 in ChimeraX, we identified Atf7IP2(C353) as a candidate for self-crosslinking. The vectors needed to express the mutant 6Xhis-Setdb1(10-108)/MBP-Atf7IP(574-666), MBP-Atf7IP(574-666), and the MBP-Atf7IP2(297-388) protein complexes were cloned and expressed as described in the Cloning and Protein expression method section.

In this experiment, the frozen pellet of each protein complex sample was thawed and cooled on ice. The samples were then resuspended in a lysis buffer containing 30mM HEPES pH 7.9, 400mM NaCl, 10% glycerol, 0.05% Tween-20 and 100 μM PMSF, sonicated and clarified by centrifugation at 10000rpm for 20 min at 4°C. The clarified supernatants were then incubated with 200µl of magnetic nickel beads or amylose beads for 2 hours at 4°C with gentle rotation. After incubation, the beads were washed three times with wash buffer (lysis buffer with 25mM imidazole) to remove nonspecifically bound proteins. Bound proteins were eluted by incubating the samples with the lysis buffer supplemented with 500mM imidazole or 20mM Maltose for 1 hour at 4°C with gentle rotation. To assess the crosslinking results, two gel samples were made for each elute, with and without BME which were distinguished through SDS-PAGE and Coomassie staining.

## Mass Spectrometry

Sample stocks containing pLK39-co-expressed MBP-Setdb1 (2-114) and His-sumo-Atf7IP (574-666) were stored in 30 mM HEPES, 400 mM NaCl, 10% glycerol, 0.05% Tween-20, 20 mM maltose, 1mM BME, pH 7.9. Prior to mass spectrometry experiments, the samples were buffer exchanged twice into 400 mM ammonium acetate (pH adjusted to 7.9 using ammonium hydroxide) and 1 mM DTT solution, using BioRad P6 size-exclusion columns. Concentration was measured using A_280_ on a NanoDrop spectrophotometer (Thermo). Native mass spectrometry (nMS) experiments were performed on Thermo Q Exactive Ultra High Mass Range (UHMR) mass spectrometer modified with a custom surface induced dissociation (SID) device^48, 49^. Sample was diluted to 500 nM and then loaded into in-house pulled glass emitters (Sutter Instruments _P-97 Tip Puller). Nano-electrospray ionization was induced by applying a voltage of 0.9 kV directly to the solution using a platinum wire inserted into the back of the spray needle. The instrument was operated in positive mode with parameters listed as follows: Capillary temperature 250°C; In-source trapping -60 V; Trapping gas 5; Resolution 3125; *m/z* scan range 1000- 15000. Collision induced dissociation (CID) experiments were performed by isolating a 500 *m/z* window from the MS1 spectrum using the quadrupole, followed by the application of an HCD voltage of 220 V (4180 eV or 3960 eV for charge states +19 and +18, respectively). Spectra were deconvolved using UniDec^50^.

## Size-exclusion chromatography Coupled with Multi-Angle Light Scattering (SEC-MALS) experiments

The molecular mass of the MBP-Setdb1/His-SUMO-Atf7IP complex was determined by Size-Exclusion Chromatography coupled with Multi-Angle Light Scattering (SEC-MALS). The experiments were conducted using a Wyatt DAWN HELEOS II multi-angle light scattering detector coupled to an ÄKTA Pure FPLC system (GE Healthcare) equipped with a Superdex 200 Increase 10/300 GL column (Cytiva). Recombinant MBP-Setdb1(2-115)/6XHis-SUMO-Atf7IP(574-666) were expressed and purified as described above. The protein complex was loaded onto the Superdex 200 Increase 10/300 GL column, pre-equilibrated with the same buffer, at a flow rate of 0.5 mL/min. The column was connected in-line with a UV detector (measuring absorbance at 280 nm), a refractive index (RI) detector (Optilab T-rEX, Wyatt Technology), and a multi-angle light scattering detector (Wyatt DAWN HELEOS II). Approximately 400 µL of the protein complex (at ∼1 mg/mL) was injected for analysis. The molecular mass of the MBP-Setdb1(2-115)/6X-His-SUMO-Atf7IP(574-666) complex was determined in real-time by measuring light scattering and refractive index. The data were analyzed using Astra software (Wyatt Technology). The molecular weight was calculated using the Zimm equation, which correlates the light scattering intensity to molecular mass without the need for a standard calibration curve. To confirm the homogeneity and oligomeric state of the complex, the elution profile from SEC was monitored for the presence of a single peak, and the molecular mass was determined from the MALS data for the corresponding peak. The resulting molecular mass was compared to the theoretical mass based on the individual components of the complex, confirming the formation of the MBP-Setdb1(2-115)/6X-His-SUMO-Atf7IP(574-666) complex. To ensure specificity and accuracy, a molecular mass standard, bovine serum albumin, BSA was run prior to running the protein complex to validate the SEC-MALS system.

## In Vivo FLAG Pull-Down and Immunoprecipitation (IP) Assays

To investigate the interaction between Flag-Atf7IP/Flag-Atf7IP(1-448)/Flag-Atf7IP(422-841)/Flag-Atf7IP(836- 1270)/ Flag-Atf7IP(Δ574-666), Myc-Atf7IP and HA-Setdb1, in vivo FLAG pull-down and co-immunoprecipitation (co-IP) assays were performed in HEK293T cells. These cells were made to transiently express these proteins in different combinations in triplicates as described in the paper. Peg BacMam plasmids encoding these genes were generated as described earlier. For co-expression conditions, cells were co-transfected with 1 µg of each protein at a 1:1:1 or 1:1 ratio. For all the single-protein conditions, 1µg of each protein was used.

The pEG BacMam vector transduction system was used as described previously^51^ to culture HEK293T cells in DMEM supplemented with 10% FBS at 37°C in a 5% CO₂ atmosphere and transiently transfect them with the plasmids using Lipofectamine 2000(Invitrogen). Briefly, the cells were seeded 24 hours before transfection. The DNA transfections were prepared with Optimem and Lipofectamine 2000 (Invitrogen) according to the protocol. Following 24h after the transfection, the transfected cells were treated 10mM sodium butyrate and incubated at 37°C in a 5% CO₂ atmosphere for another 8-24 hours. At the 24-hour mark post-transfection, the media was discarded and, the cells were collected with 1mL of Tris-buffered saline (TBS) and harvested by centrifugation at 1500g for 5 min at 4°C. The supernatants were discarded, and the cell pellets were then flash-frozen in liquid nitrogen and stored at −80°C until ready to use.

Per sample, cells were lysed in 1mL of lysis buffer containing 30mM HEPES pH 7.9, 400mM NaCl, 10% glycerol, 0.05% Tween-20, 2mM BME, 50U of high-salt resistant Benzonase (Sigma Aldrich), 1 mM PMSF, and protease inhibitor cocktail (Sigma-Aldrich) on ice for 1 hour. Lysates were clarified by centrifugation at 13,200 rpm for 10 min at 4°C.The clarified lysate was incubated with 200µl of MonoRab™ Anti-DYKDDDDK Magnetic Beads (Genscript) at 4°C for 1 hour with gentle rotation. The beads were washed three times with lysis buffer to reduce nonspecific binding. Bound proteins were eluted by incubating the beads with lysis buffer supplemented with 200 µg/mL FLAG peptide (Sigma-Aldrich) for 1 hour at 4°C with gentle rotation.

Eluted proteins from these FLAG pull-down assays were separated by SDS-PAGE on a 12% gel and transferred to a PVDF membrane (Millipore). The membranes were blocked with 5% non-fat dry milk in TBS-T (Tris-buffered saline with 0.1% Tween-20)] for 1 hour at room temperature, washed thrice with TBS-T and probed overnight at 4°C with the following primary antibodies: anti-FLAG (F1804, Sigma-Aldrich, 1:5000), anti-HA (Invitrogen, 1:4000), and anti-Myc (9B11, Cell Signaling Technology, 1:5000). After washing, the membranes were incubated with HRP-conjugated secondary antibodies (1:10000) for 1 hour at room temperature. Protein bands were visualized using an enhanced chemiluminescence (ECL) detection system (Thermo Fisher Scientific), and images were captured using the Bio-Rad ChemiDoc imaging system.

## *In vitro* CRM1-binding assays

Each 200 μL binding assay reaction contains 20 μL of glutathione Sepharose beads (Cytiva) with 0.5 μM GST- Setdb1 protein immobilized and 1.5 μM MBP-Atf71P in TB Buffer (20mM HEPES 7.4, 110mM potassium acetate, 2mM magnesium acetate, 10% glycerol, 2mM DTT, 0.005% Tween20). Binding assay reactions were similarly assembled using 0.5 μM GST protein and GST-X11L2^NES^ as controls. The samples were rotated at 4°C for 30 minutes before addition of 1.5 μM of purified hCRM1 with or without 4.5 μM Ran^GTP^ followed by mixing for another 30 minutes at room temperature. After extensive washing, the bound and unbound proteins were separated and visualized by Coomassie stained SDS-PAGE.

