## Supplement for "Setdb1 and Atf7IP form a hetero-trimeric complex that blocks Setdb1 nuclear export"

### Figure S1

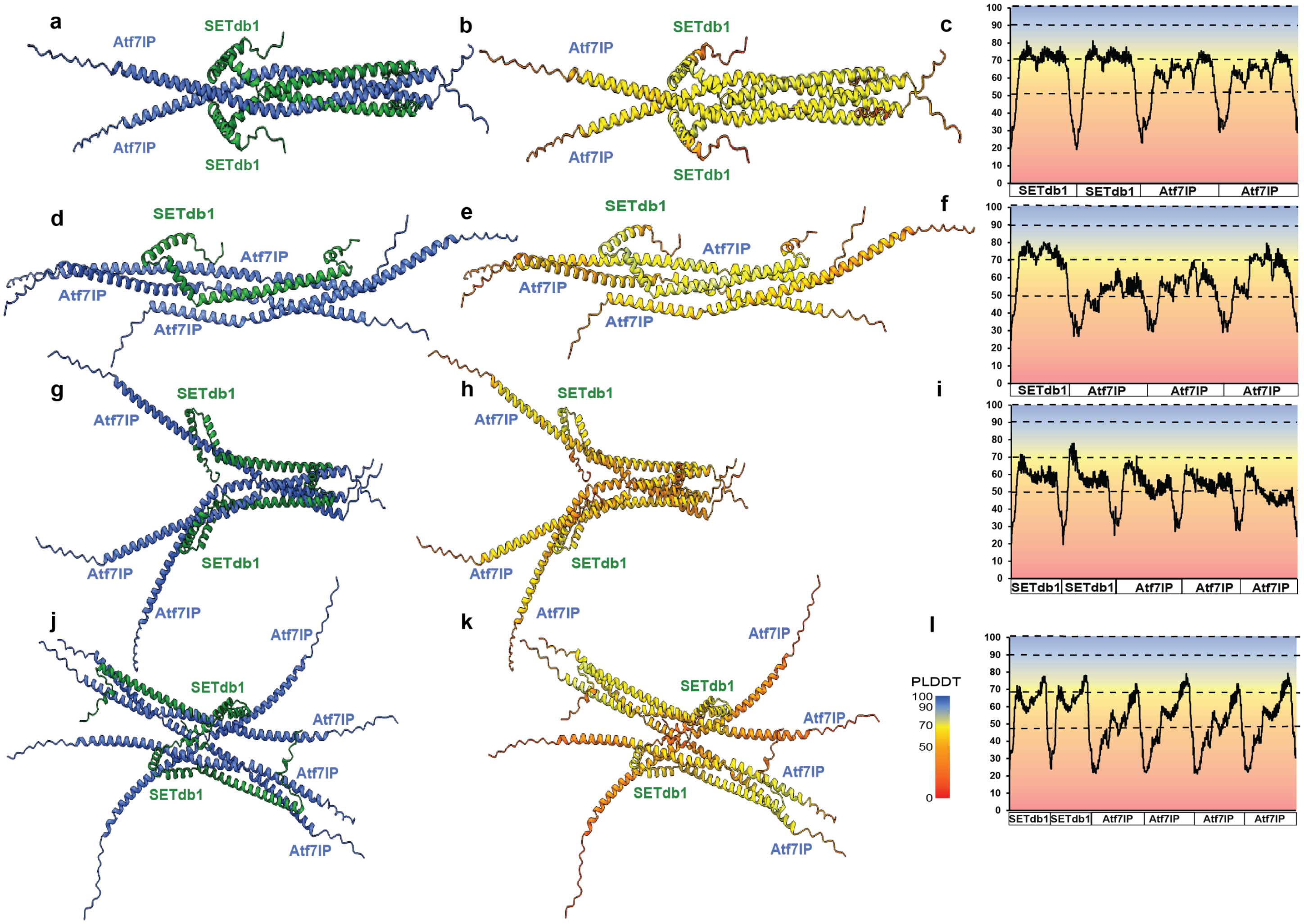


**Figure S1: The low-confidence Alphafold2 models of the minimal SETdb1/Atf7IP complex with different stoichiometries.** These models represent the different stoichiometries tested for the SETdb1/Atf7IP minimal interaction, other than the 1:2 interaction; a) 2SEtdb1:2At7IP d) 1SETdb1: 3Atf7IP g) 2SETdb1:3Atf7IP j) 2SETdb1:4Atf7IP The right (b-c, e-f, h-i, k-l) of each panel contains the pLDDT-colored model along with its pLDDT plot predicted for each stoichiometry.

#### Figure S2

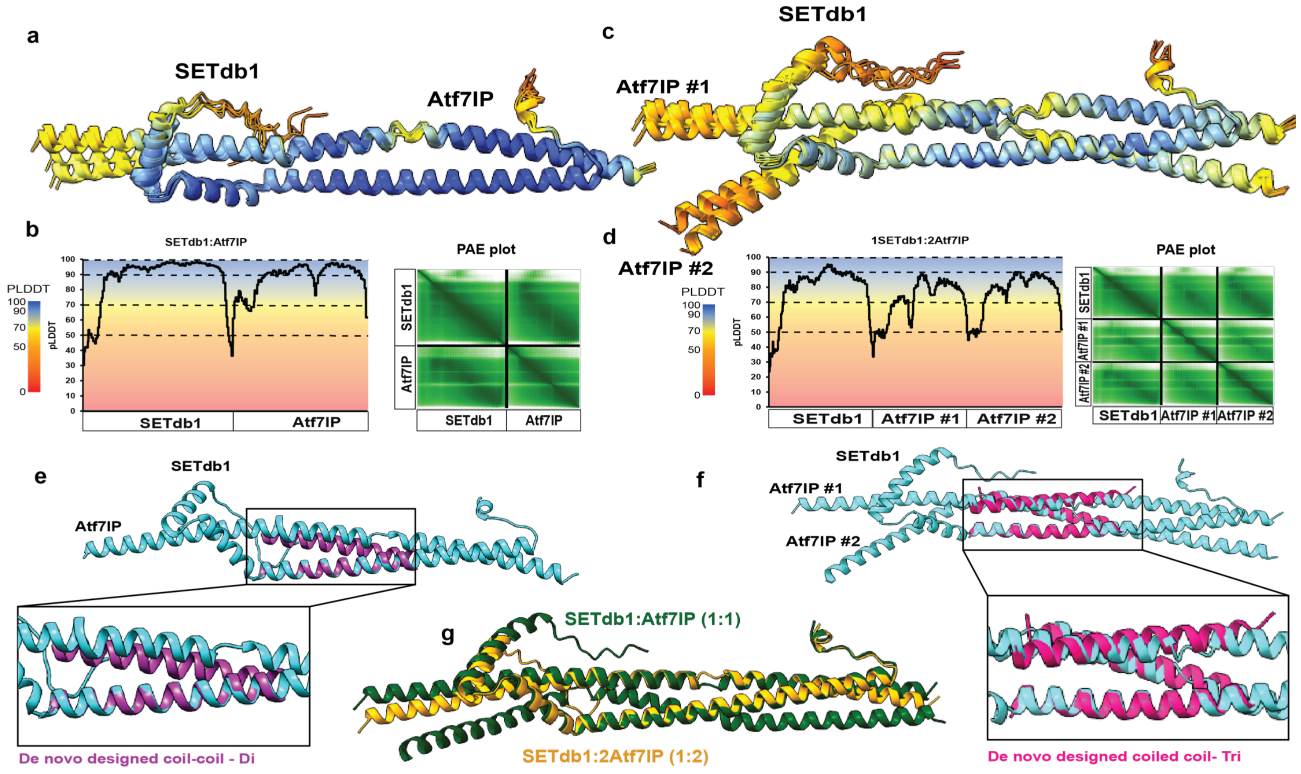


**Figure S2: SETdb1/Atf7IP 1:1 AF2 model has a slightly higher pLDDT and PAE scores than its 1:2 AF2 model.** a,c) Alignment of five top ranked models of the SETdb1(2-115)/Atf7IP(574-666) 1:1 and 1:2 models colored by pLDDT scoring. b) pLDDT plot and PAE plot of SETdb1/Atf7IP 1:1 highest ranked model d) pLDDT plot and PAE plot of SETdb1/Atf7IP 1:2 highest ranked model e,f) Superimposed idealized coiled-coils (pdb: 4dzl and 4dzm) with the dimeric and hetrotrimeric models of the SETdb1/Atf7IP minimal complex respectively. g) Superimposed model of the 1:1 and 1:2 SETdb1/Atf7IP minimal complexes.

#### Figure S3

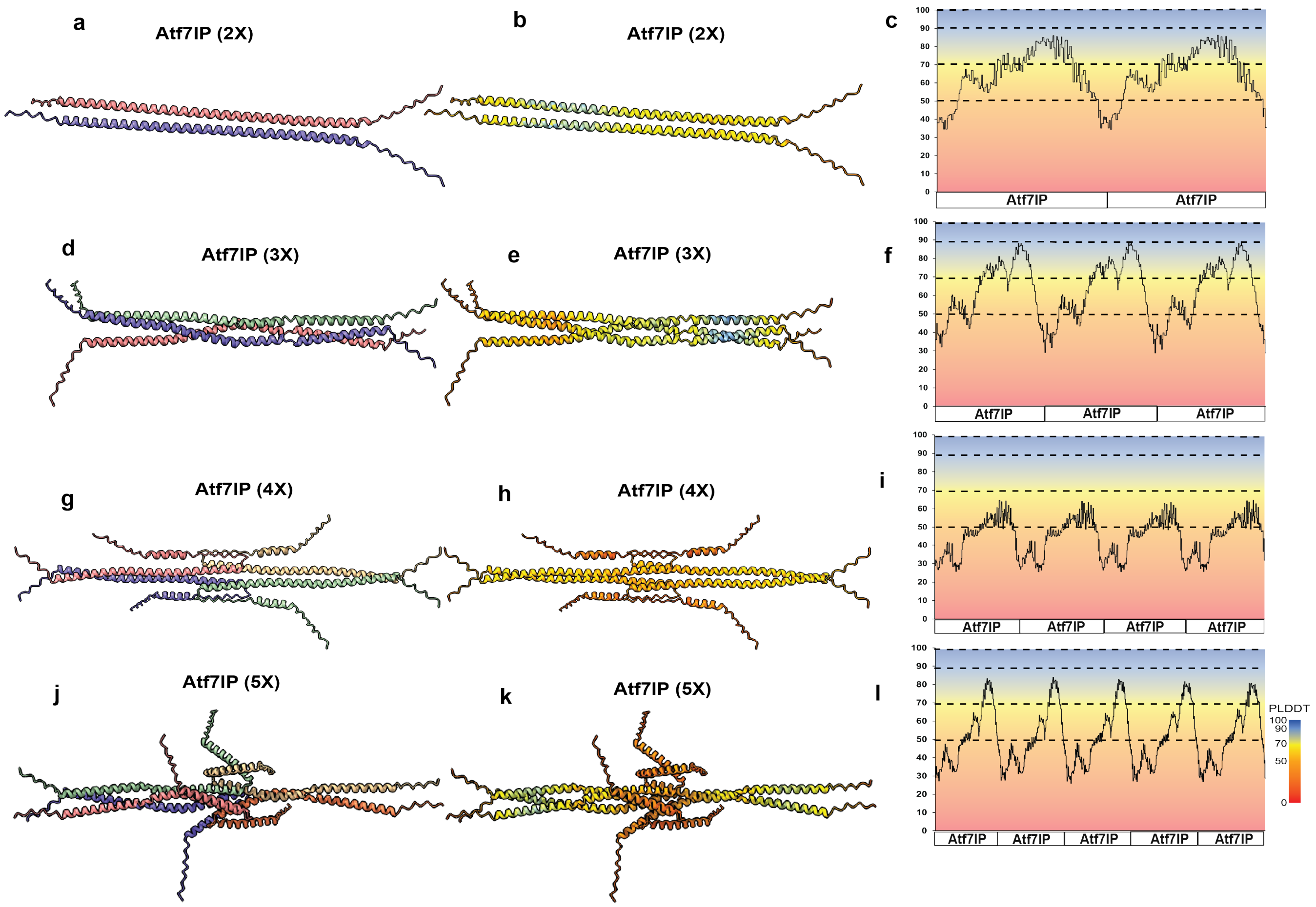


**Figure S3 Alphafold2 models of Atf7IP(574-666) fragments with itself do not support self-dimerization.** These models represent the different stoichiometries tested for the Atf7IP self-oligomerization; a) 2Atf7IP d) 3Atf7IP g) 4Atf7IP j) 5Atf7IP The right (b-c, e-f, h-i, k-l) of each panel contains the pLDDT-colored model predicted for each stoichiometry with its corresponding pLDDT plot.

### Figure S4

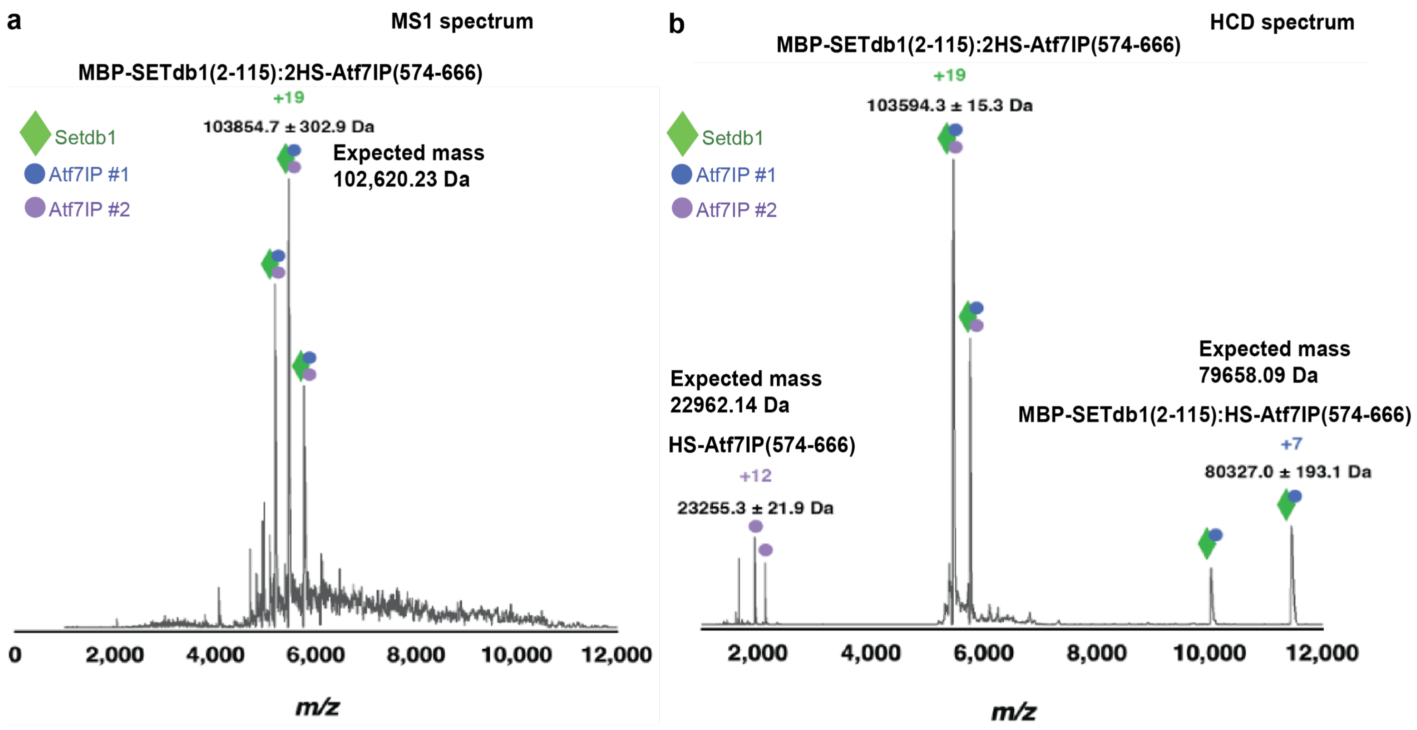


**Figure S4 Native mass spectrometry validates heterotrimeric stoichiometry of the MBP-SETdb1(2-115)/HS-Atf7IP(574-666) complex.**
a) Raw MS1 spectrum obtained on UHMR showing charge state distribution of possible 1 MBP-SETDB1(2-115): 2(6X-SUMO-Atf7IP(574-666) complex. b) HCD 220 spectrum following isolation of the 5300-5800 m/z. Most abundant charge states in each distribution are labeled. The values reported are from an average of three measurements. Representative spectrum from a single measurement shown.

### Figure S5

```
Atf7IP          MDSLEEPQKKVFKARKTMRVSDRQQLEAVYKVKEELLKTDVKLLNGNHENGDLDPTSPLENMDYIKDKEEVNGIEEICFDPEGSKAEWKETPCILSVNVKNKQDDDLNCEPLSPHNITPE 120
Atf7IP2         ------------------------------------------------------------------------------------------------------------------------ 0
Atf7IP          PVSKLPAEPVSGDPAPGDLDAGDPASGVLASGDSTSGDPTSSEPSSSDAASGDATSGDAPSGDVSPGDATSGDATADDLSSGDPTSSDPIPGEPVPVEPISGDCAADDIASSEITSVDLA 240
Atf7IP2         ------------------------------------------------------------------------------MASPDRSKRKILKA--KKTMPL--SCRKQVEMLNKSRNVEA- 37
                                                                                              ::* * :. . : .    . *:  .*  :    .:  .*:
Atf7IP          SGAPASTDPASDDLASGDLSSSELASDDLATGELASDELTSESTFDRTFEPKSVPVCEPVPEIDNIEPSSNKDDDFLEKNGADEKLEQIQSKDSLDEKNK--ADNNIDANEETLE---TD 355
Atf7IP2         -----LKTAIGSNVPSG----------------------------NQSFSPSVIT---RTTEITKCSPSENGASSLDS------------NKNSISEKSKVFSQNCIKPVEEIVHSETKL 109
                      .   ..:: **                            :::*.*. :     . ** : .**.*  ..: .            .*:*:.**.*  ::* *.  ** :.   .
Atf7IP          DTTICSDRPPENEKKVEEDIITELALGEDAISSSMEIDQGEKNEDETSADLVETINENVIEDNKSE---------NILENT------------DSMETDEIIPILEKL-------APSED 447
Atf7IP2         EQVVCSYQKPSRTTESPSRVFTEEAK--DSLNT------SE-NDSEHQTN----VTRSLFEHEGACSLKSSCCPPSVLSGVVQMPESTVTSTVGDKKTDQMVFHLETNSNSESHDKRQSD 216
                : .:** : *.. .:  . ::** *   *::.:      .* *:.* .::    :...::*.: :          .:*...            .. :**:::  **.          ..*
Atf7IP          ELTCFSKTSLLPIDETNPDLEEKMESSFGSPSKQESSESLPKEAFLVLSDEEDISGEKDESEVISQNETCSPAEVESNEKDNKPEEEEQVIHEDDERPSEKNEFSRRKRSKSEDMDNVQS 567
Atf7IP2         NILCSEDSGFVPVEKTPN-LVNSVT------------------------------S-NNCADDILKTDECSRTSISNCESADSTWQ--SSLD------TNNNSHYQKKRMFSENEENVKR 296
                :: * ..:.::*:::*   * :.:                               . :: :: * :.: ** :.:.. *. :.  :  . :.      :::*.. ::**  **: :**:
Atf7IP          KRRRYMEEEYEAEFQVKITAKGDINQKLQKVIQWLLEEKLCALQCAVFDKTLAELKTRVEKIECNKRHKTVLTELQAKIARLTKRFEAAKEDLKKRHEHPPNPPVSPGKTVNDVNSNNNM 687
Atf7IP2         ---MKTSEQINENICVS----LERQTAFLEQVRHLIQQEIYSINYELFDKKLKELNQRIGKTECRNKHEGIADKLLAKIAKLQRRIKTVLLF--QRNCLKPNMLSSNGAS--KVANS--- 402
                      .*: : :: *.     : :  : : :: *::::: :::  :***.* **: *: * **.::*: :  :* ****:* :*:::.     :*:   **   * * :  .* ..
Atf7IP          SYRNAGTVRQMLESKRNVSESAPPSFQTPVNTVSSTNLVTPPAVVSSQPKLQTPVTSGSLTATSVLPAPNTATVVATTQVPSGNPQPTISLQPLPVILHVPVAVSSQPQLLQSHPGTLVT 807
Atf7IP2         --------EAMILD-KNL-----ES-------------------------VNSPIEKSSVNYE-----PS---------NPSEK--------------------GSKKINLSSDQNKSVS 449
                        . *: . :*:      *                         :::*: ..*:.       *.          ** :                    .*:   *.*. .. *:
Atf7IP          NQPSGNVEFISVQSPPTVSGLTKNPVSLPSLPNPTKPNNVPSVPSPSIQRNPTASAAPLGTTLAVQAVPTAHSIVQATRTSLPTVGPSGLYSPSTNRGPIQMKIPISAFSTSSAAEQNSN 927
Atf7IP2         ESNNDDVMLISVESPNLTTPITSNPTDTRKI-----------------------------------------------------------------------------------TSGNSS 486
                :. ..:* :***:**  .: :*.**..  .:                                                                                   :. **.
Atf7IP          TTPRIENQTNKTIDASVSKKAADSTSQCGKATGSDSSGVIDLTMDDEESGASQDPKKLNHTPVSTMSSSQPVSRPLQPIQPAPPLQPSGVPTSGPSQTTIHLLPTAPTTVNVTHRPVTQV 1047
Atf7IP2         NSPNAEVM----------------------AVQKKLDSIIDLTKE----------------------------------------------------------------------GLSNC 514
                .:*. *                        *. .. ..:**** :                                                                       :::
Atf7IP          TTRLPVPRAPANHQVVYTTLPAPPAQAPLRGTVMQAPAVRQVNPQNSVTVRVPQTTTYVVNNGLTLGSTGPQLTVHHRPPQVHTEPPRPVHPAPLPEAPQPQRLPPEAASTSLPQKPHLK 1167
Atf7IP2         NTESPV--------------------SPLESHSKAASNSKETTPLAQ--------------------------NAVQVPESFEHLPPLPEPPAPLPELV------DKTRDTLPPQKPELK 582
                .*. **                    :**..    *   ::..*  .                          .. : * ...  ** *  ******         :: .*  ****.**
Atf7IP          LARVQSQNGIVLSWSVLEVDRSCATVDSYHLYAYHEEPSATVPSQWKKIGEVKALPLPMACTLTQFVSGSKYYFAVRAKDIYGRFGPFCDPQSTDVISSTQSS  1270
Atf7IP2         VKRVFRPNGIALTWNITKINPKCAPVESYHLFLCHENSNN--KLIWKKIGEIKALPLPMACTLSQFLASNRYYFTVQSKDIFGRYGPFCDIKSIPGFSENLT-  682
                : **   ***.*:*.: ::: .** *:****:  **: .      ******:***********:**::..:***:*::***:**:***** :*   :*.. :
```

**Figure S5 Clustal-O sequence alignment of Atf7IP and Atf7IP2.**

The red box highlights the coiled-coil regions of Atf7IP and Atf7IP2. Abbreviations: * = identical residues, : = strongly similar residues, . = moderately similar residues

### Figure S6

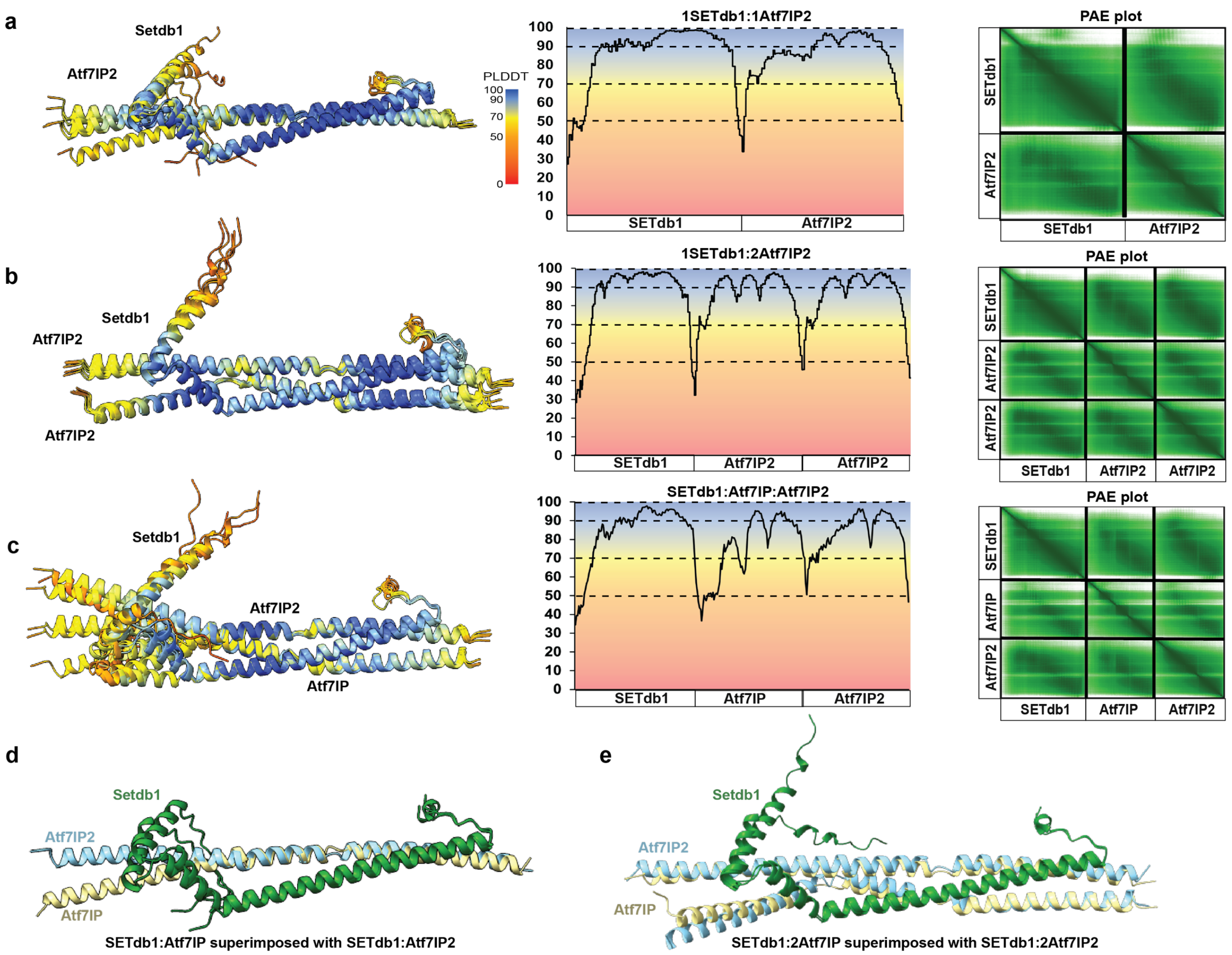


**Figure S6 : Alphafold2 models of the minimal 1:1 and 1:2 SETdb1/Atf7IP2 complexes are similar to the m.** a-b) Alignment of five top ranked models of the SETdb1(2-115)/Atf7IP2(297-388) 1:1 and 1:2 models colored by pLDDT scoring with their corresponding pLDDT plot and PAE plot of the highest ranked model c-d) Superimposed models of the 1:1 and 1:2 SETdb1/Atf7IP2 with the SETdb1/Atf7IP minimal complexes.

### Figure S7

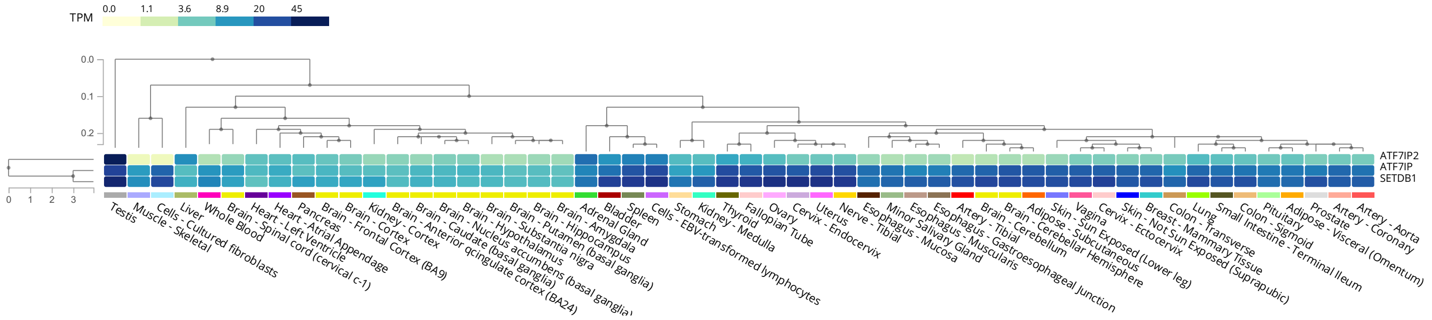


**Figure S7: Differential expression of Setdb1, Atf7IP and Atf7IP2 in human tissues.** Plot was generated using data from GTEx portal. TPM; transcripts per million.
